# Spotting genome-wide pigmentation variation in a brown trout admixture context

**DOI:** 10.1101/2020.07.23.217109

**Authors:** T. Valette, M. Leitwein, J.-M. Lascaux, E. Desmarais, P. Berrebi, B. Guinand

## Abstract

Variation in body pigmentation attracted fish biologists for a while, but high-throughput genomic studies investigating its molecular basis remain limited to few species and associated conservation issues ignored. Using 75,684 SNPs, we explored the genomic basis of pigmentation pattern variation among individuals of the Atlantic and Mediterranean clades of the brown trout (*Salmo trutta*), a polytypic species in which Atlantic hatchery individuals are commonly used to supplement local wild populations. Using redundancy analyses and genome-wide association studies, a set of 337 independent “colour patterning loci” (CPLs) significantly associated with pigmentation traits such as the number of red and black spots on flanks, or the presence of a black spot on the pre-opercular bone was identified. CPLs map onto 35 out of 40 brown trout linkage groups indicating a polygenic basis to pigmentation patterns. They are mostly located in coding regions (43.4%) of 223 candidate genes, and correspond to GO-terms known to be involved in pigmentation (e.g. calcium and ion-binding, cell adhesion). Annotated candidates include genes with known pigmentation effects (e.g. *SOX10*, *PEML, SLC45A2*), but also the Gap-junction Δ2 (*GJD2*) gene already shown differentially expressed in trout skin. Patterns of admixture were found significantly distinct when using either the full SNP data set or the set of CPLs, indicating that pigmentation patterns accessible to practitioners are not a reliable proxy of genome-wide admixture. Consequences for management are discussed.

## 1. INTRODUCTION

The study of pigmentation has a peculiar place in modern biology as colour and its patterning play important roles in natural, artificial and sexual selection (Cieslak, Reissmann, Hofreiter, & Ludwig, 2011; Cuthill et al., 2017). The biology and evolution of animal colouration largely benefited over the past few years in the development of high-throughput sequencing techniques (San José & Roulin, 2017; Orteu & Jiggins, 2020). While supergenes and large effect mutations have been shown to support pigmentation patterns in many organisms (Orteu & Jiggins, 2020), genomic studies also generated a huge literature that underscored their polygenic basis (e.g. skin and hair colour in human: Crawford et al., 2017; Pavan & Sturm, 2019; structural colour variation or eyespot numbers in butterflies: Brien et al., 2019; Rivera-Colón, Westermann, van Belleghem, Monteiro, & Papa, 2020). Recent research showed that pigmentation variation may result from effects propagated during development by loci belonging to regulatory networks to hub or master genes often sufficient to explain most of the causal variation with colour expression and/or patterning (e.g. Arnould et al., 2013; Ordway, Hancuch, Johnson, Wiliams, & Rebeiz, 2014; Ding et al., 2020, Orteu & Jiggins, 2020).

If early genetic and molecular studies were interested in colour and pigmentation patterns in fish – e.g. sex-linked colour variation (Kottler & Schartl, 2018) – the issue of body pigmentation gained interest with the establishment of zebrafish and medaka as model species (Parichy, 2006; Takeda & Shimada, 2010; Singh & Nüsslein-Völlard, 2015; Nüsslein-Völlard & Singh, 2017) and from comparisons with higher vertebrates (e.g. Kelsh, Harris, Colanesi, & Erickson, 2009). Pigment cells (i.e. chromatophores; mainly melanophores, iridophores, leucophores and xanthophores) are distributed in the hypo- and the epidermis in fish, and mutational or other analyses allowed for increased understanding in the mechanisms of pigment cell fates during development and their resulting distribution (Kelsh et al., 2004, 2009; Kimura et al., 2014; Eom, Bain, Patterson, Grout, & Parichy, 2015; Nüsslein-Völlard & Singh, 2017; Parichy & Spiewak, 2015; Singh & Nüsslein-Völlard, 2015; Salis et al., 2018; Volkening 2020). Studies now extend to other fish species (Maan & Sfec, 2013; Irion & Nüsslein-Völlard, 2019) and to a large array of eco-evolutionary questions regarding the involvement of genes or regulatory pathways in fish pigmentation, its epistatic and pleiotropic nature, its modularity and its control, its role in speciation as well as the impact of whole-genome duplication that promoted the diversification of pigment cell lineages (Hultman, Bahary, Zon, & Johnson, 2007; Miller et al., 2007; Braasch, Brunet, Volff, & Schartl, 2009; Roberts, Ser & Kocher, 2009; Albertson et al., 2014; Santos et al. 2014; Ceinos, Guillot, Kelsh, Cerdá-Reveter, & Rotlland, 2015; Yong, Peichel, & McKinnon, 2015; Kimura, Takehana, & Naruse, 2017; Roberts, Moore, & Kocher, 2017; Lorin, Brunet, Laudet, & Volff, 2018; Kratochwil et al., 2018; Nagao et al., 2018; Cal et al., 2019; Lewis et al., 2019; Liang, Gerwin, Meyer, & Kratochwil, 2020). Studies also engaged fish research in high-throughput genomic approaches of pigmentation variation (Tripathi et al., 2009; Greenwood et al., 2011; Malek, Boughman, Dworkin, & Peichel, 2012; O’Quin, Drilea, Conte, & Kocher, 2013; Henning, Jones, Franchini, & Meyer, 2013; Albertson et al., 2014; Henning, Lee, Franchini, & Meyer, 2014; Xu et al., 2014; Bian et al., 2016; Zhu et al., 2016; Roberts et al. 2017; Kon et al., 2020). However, in spite of this impressive research, the genomics of pigmentation variation remain poorly investigated in fish compared to other phenotypic traits (Peichel & Marques, 2017). Furthermore, links between pigmentation and conservation genomic issues remain absent (but see Boulding et al., 2008) while this topic emerges in, e.g., birds, notably regarding the impacts of admixture (Toews et al., 2016; Hanna, Dumbacher, Bowie, Henderson, & Wall, 2018; Billerman, Cicero, Bowie, & Carling, 2019).

Salmonids represent a large fish family with a diverse and complex body pigmentation going from continuous colour to spotty, marbled, blotchy and striped patterns. If whole body colouration or specific coloured elements have a genetic basis (e.g. Blanc, Poisson, & Vibert, 1982; Skaala & Jørstad, 1988; Blanc, Chevassus, & Krieg, 1994; Blanc, Poisson, & Quillet, 2006; Boulding et al., 2008; Colihueque, 2010; Nilsson et al., 2016), another part is plastic (Colihueque, 2010; Westley, Stanley, & Fleming, 2013; Jørgensen et al., 2018). Colour patterns support differences in coping styles (Kittilsen et al., 2009; Brännäs et al., 2016) or in the physiological adjustments necessary to avoid predation and match with environmental variability (e.g. Miyamoto, 2016; Jacquin et al., 2017; Zastavniouk, Weir, & Fraser, 2017). In salmonids, pigmentation and colour are also known to interact with social hierarchies, influence mate choice and affect fitness (e.g. O’Connor, Metcalfe, & Taylor, 2000; Wedekind, Jacob, Evanno, Nusslé, & Müller, 2008; Marie-Orléach et al., 2014; Watt, Swanson, Miller, Chen, & May, 2017, Auld, Noakes, & Banks, 2019). However, the genomic basis of their pigmentation patterns remains largely unexplored, with limited insights coming from quantitative trait loci (QTL) (Boulding et al., 2008) or few gene expression studies (Sivka, Snoj, Palandaćič, & Sušnik Bajec, 2013; Djurdjevič, Furmanek, Miyazawa, & Sušnik Bajec, 2019). Furthermore, as pigmentation patterns may reflect admixture in salmonids (Largiadèr & Scholl, 1996; Mezzera, Largiadèr, & Scholl, 1997; Aparicio, García-Berthou, Araguas, Martinez, & Garcia-Marin, 2005; Miyazawa, Okamoto, & Kondo, 2010; Kirczuk & Domagała; 2012, Kocaba, Kutluyer, & Başçinar, 2018), it appears necessary to improve knowledge on the genotype-phenotype association supporting observed pigmentation patterns. Indeed, these patterns may inform on individual origin and use to improve management knowledge, decisions or policies (Largiadèr & Scholl, 1996; Poteaux & Berrebi, 1997; Delling, Crivelli, Rubin, & Berrebi, 2000; Aparicio et al., 2005; Kocabaş et al., 2011; Marin, Coon, & Fraser, 2017; Duchi, 2018; Lorenzoni et al., 2019). It is thus important to evaluate if loci supporting phenotypic divergence for pigmentation patterning reflect genome-wide admixture.

In this study, we take advantage of former genomic knowledge established in the brown trout (Leitwein, Gagnaire, Desmarais, Berrebi, & Guinand, 2018) to search for the loci responsible for body pigmentation variation among populations of two clades recognised in this species (Atlantic and Mediterranean; Bernatchez, 2001; Sanz, 2018). Then, we investigate if those loci could be a relevant proxy to monitor hybridization and genome-wide admixture among individuals of these clades.

## 2. MATERIALS AND METHODS

### 2.1 Brown trout’s genomics data

Double digested restriction site-associated DNA sequencing (ddRADseq) data used in this study are from Leitwein et al. (2018). The data set consists in 75,684 genome-wide SNPs (40,519 RAD-loci or haplotypes) for 112 trout of hatchery and wild caught origins. Eighty-two wild caught individuals were fished in the headwaters of three rivers within the Mediterranean Orb River catchment (France) (Gravezon, Mare and Upper Orb rivers; Leitwein et al., 2018). This catchment has been seeded by hatchery fish of both Atlantic and Mediterranean origins for decades. This stopped in 2004, but unintentional releases due to flooding of private hatcheries are documented (Leitwein et al., 2018). Analyses showed that these wild caught individuals consisted in hatchery individuals of both Mediterranean and Atlantic origins, F1’s, F2’s, backcrossed individuals, and ‘pure’ natural Mediterranean fish (Leitwein et al., 2018). ‘Early’ and ‘late’ backcrossed individuals correspond to Mediterranean wild-caught fish that contained distinct distributions of Atlantic ancestry tracts in their genome. Thirty individuals of the domestic Atlantic (*N* = 15) and Mediterranean (*N* = 15) strains were also included in the analysis. Mediterranean hatchery fish have been randomly sampled in a local strain formerly established using mature trout from the Gravezon River by the Fédération de Pêche de l’Hérault in 2004. Atlantic hatchery fish originated from the Cauterets hatchery that maintains an Atlantic strain distributed worldwide (Bohling, Haffray, & Berrebi, 2016).

### 2.2 Phenotypic data

Acquisition of phenotypic data followed Lascaux (1996) and is used by numerous angler associations and management authorities in France. The set of variables considered in this protocol is listed in Table S1 and illustrated on a picture in Fig. S1. Data were recorded from individual photographs of the 112 fish considered for genomic analyses. Photographs were taken at fishing on slightly anesthetized trout with eugenol. A camera Canon® EOS 1000D was used. After recovery, fish were released in the wild or in hatchery tanks. Quantitative variables (*N* =19) and semi-quantitative variables (*N* =11) were recorded by visual examination of photographs of the left flank of each individual fish. Quantitative variables refer to punctuation patterns of brown trout (e.g. number and diameter of spots) and are recognized important for this species (Blanc et al., 1982, 1994). Semi-quantitative variables refer to ‘ornamental’ appearance patterns (e.g. parr marks, fringes on fins, white ring around spots) that are important to practitioners or anglers. Each quantitative punctuation variable was measured independently. Ornamental variables have been coded by modalities, either present/absent [1, 2] or, e.g., absent/partial/complete [1, 2, 3] (Tables 1 & S1). This set of variables avoided focusing only on peculiar attributes of the pigmentation patterns. Other pigmentation traits that can be also important for trout (e.g. background colour, brightness, or reflectance; Colihueque, 2010) were not considered in this study. Phenotypic data and trout pictures will be deposited on a public repository. Eleven colour patterning variables were found not significantly correlated (Table S2) and retained for subsequent analysis (Table 1).

**Table 1:** Uncorrelated pigmentation variables retained to build the forward selection model in RDA, and also used as a basis to multi-GWAS modelling. The full list of variables is reported in Table S1; correlation among variables in Table S2.

| Phenotypic variables | Description | Nature* (coding) |
| --- | --- | --- |
| *N. PR. Tot* | Total number of red spots over flanks | Q |
| *N. PN. Tot.* | Total number of black spots over flanks | Q |
| *Diam. PN* | Mean diameter of black spots | Q |
| *Oc. PN* | White ring around black spots | SQ (1= no ring; 2 = attenuated/thin ring, 3 = marked/thick ring) |
| *Fr. D* | Fringe on the dorsal fin | SQ (1= no fringe; 2 = white fringe, 3 = black & white fringe) |
| *Fr. An* | Fringe on the anal fin | SQ (1= no fringe; 2 = white fringe, 3 = black & white fringe) |
| *Fr. P* | Fringe on the pelvic fins | SQ (1= no fringe; 2 = white fringe, 3 = black & white fringe) |
| *L. Lat* | Visibility of the lateral line | SQ (1 = not visible; 2 = partially visible; 3 = fully visible) |
| *Parr* | Presence of parr marks | SQ (1 = absence; 2 = presence) |
| *Macrost.* | Macrostigma spot (large black dot on the pre-opercular bone) | SQ (1 = absence; 2 = partially visible/diluted; 3 = fully visible) |
| *Zeb* | Zebras on flank | SQ (1 = absence; 2 = presence) |

\* : Q = quantitative variable; SQ = semi-quantitative variable (coding in brackets)

**Table 2:**
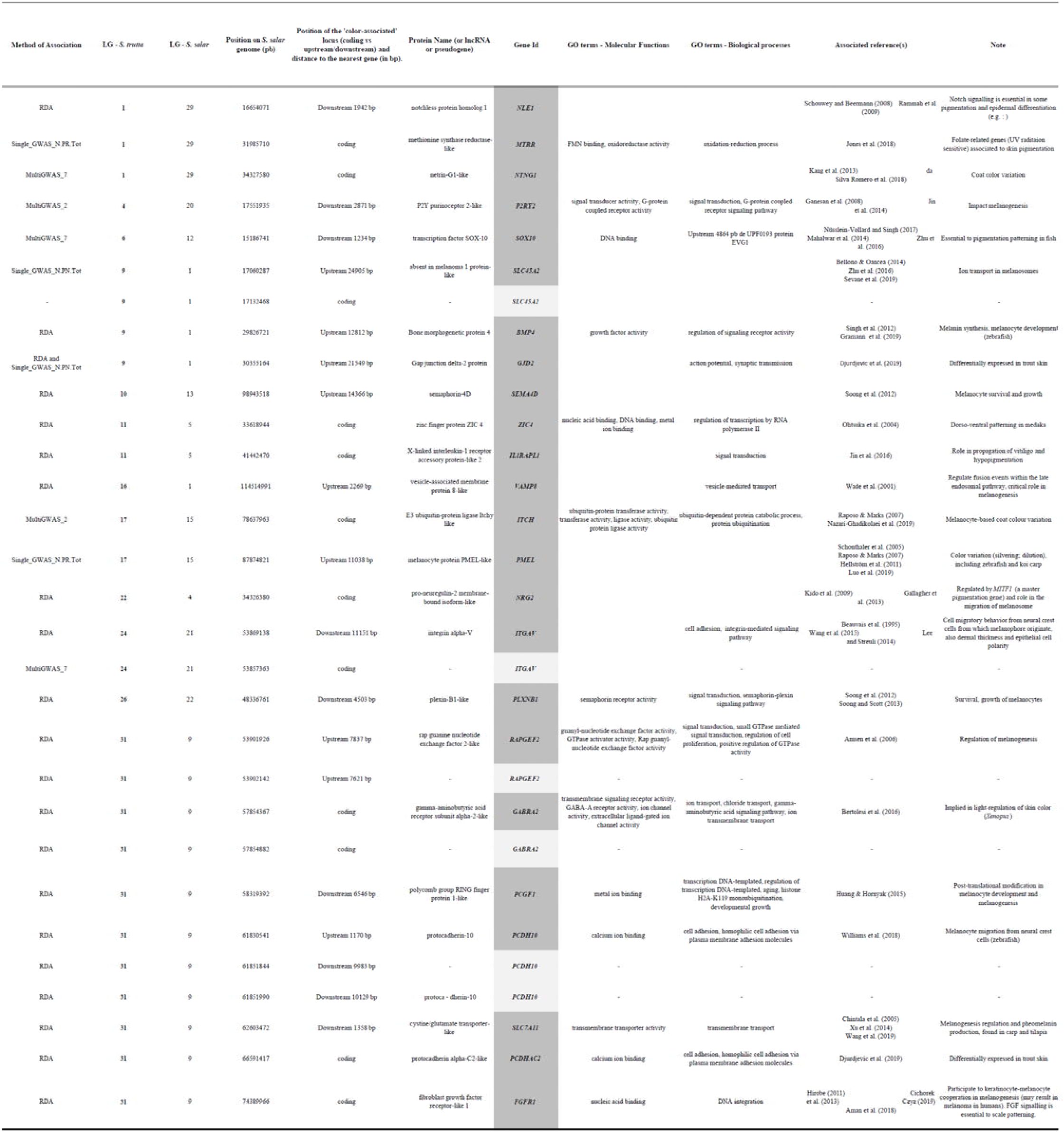
List of the most indicative pigmentation- or colour-related genes associated to some CPLs (*N* = 24; 30 distinct loci) found in this study. The method of association that detected them is reported, together with their locations in the Atlantic salmon (*Salmo salar*) genome (Lien et al., 2016) and brown trout (*S. trutta*) high density linkage map of Leitwein et al. (2017). Their position in coding sequence or in a 25kb window downstream or upstream the gene is reported. Significant GO-terms are reported. For each gene, relevant references with link to colour and/or skin features and pigmentation are provided. The full list of CPLs and other relevant information is given in Table S4.

### 2.3 Genotype-phenotype association

The evaluation of the genotype-phenotype association between SNPs and pigmentation/colour traits was performed using three distinct approaches. Hereafter, we coined a ‘colour patterning locus’ (CPL) any SNP or RAD-locus found significantly associated with a colour variable considered in this study. A CPL is not a ‘pigmentation locus’ (or gene, if any; see Lorin et al., 2018), but a locus that participate to produce pigmentation and colour information.

*Redundancy analysis (RDA)* - We first used RDA to investigate any association between genomic and colour trait data. RDA is a constrained ordination method in which only the variation of response variables explained by a set of explanatory variables is displayed and analysed (Legendre & Legendre, 2012). RDA was performed with the remaining 11 uncorrelated colour variables as the explanatory and the SNPs as the response variables. Missing genomic data were imputed by the most commonly observed genotype. Missing phenotypic data were imputed by the mean of observed trait for quantitative data, and the most commonly observed phenotype for semi-quantitative data. We classified SNPs as showing statistically significant association with individual pigmentation/ornamental traits when they loaded with more than 2.5 standard deviations (S.D.) from the mean.

A forward model selection was used to select for the relevant phenotypic variables structuring the RDA (Blanchet, Legendre, & Borcard, 2008). In constrained ordination methods like RDA, increasing the number of explanatory variables becomes similar to unconstrained ordination method (e.g. principal component analysis) as the percentage of variation explained increases when considering more explanatory variables, while some of them add no relevant information. Models were defined, first including the eleven uncorrelated response variables, then reducing this number. The Aikake Information Criterion was computed in each case to select the most appropriate model (i.e. minimizing deviance). Permutation tests (*N* = 999) were performed by permuting individuals in each model. This procedure was first established for the RDA itself, then for each successive RDA axis to investigate if observed patterns carried significant association between relevant pigmentation traits and SNPs. Analyses were performed with the *vegan* package (https://cran.r-project.org/web/packages/vegan/index.html). Once relevant colour variables were identified in models, each significant SNP was associated to the pigmentation or ornamental trait it was the most significantly correlated.

*Genome-wide association studies (GWAS)* –A single-trait association was first considered by fitting each colour variable with SNPs by linear regression. We used a penalized maximum-likelihood least absolute shrinkage and selection operator (LASSO) model to select for SNPs implied in each trait association, then solving:

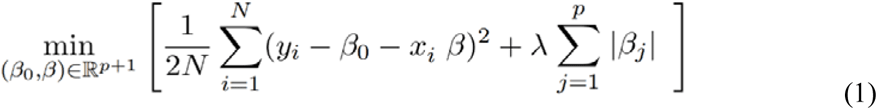

in which *Y* ∈ ℝ represents the response colour variable, *X* ∈ *ℝ^p^* a vector of predictor variables (i.e. SNPs), λ the penalty parameter, β*_0_* the y-intercept of multiple linear regression, and β *∈ ℝ^p^* a vector of β*_j_* coefficients (Friedman, Hastie, & Tibshirani, 2010). This vector of β*_j_* coefficients represents the effect size β*_j_* of the *j*^th^ SNP conditional on the effects of all other SNPs. The penalized term λ shrinks the regression coefficient towards zero, keeping only a small number of SNPs with large effects in the model. A cyclical coordinate descent procedure was retained for model selection (Friedman et al., 2010). The retained model was determined by cross-validation. Log(λ) was estimated by minimizing the mean quadratic error. The number of positive β*_j_* coefficients was estimated from log(λ), with each β*_j_* coefficient associated to a suite of SNPs considered as involved in the association. Analyses were performed with the *glmnet* package (https://cran.r-project.org/package=glmnet).

A multi-trait GWAS was also implemented. We used the MultiPhen package (O’Reilly et al., 2012) to test for the linear combination of phenotypes most associated with the genotypes at each SNP. Such one approach may potentially capture effects hidden to single phenotype GWAS. It performs a ‘reversed regression’, with multiple phenotype predictors and genetic variant as outcome (i.e. *G* SNPs *X_(i)_ = {X_(i1)_,…, X_(iG)_}* are explained by *K* pigmentation variables *Y_(i)_ = {Y_(i1)_,…, Y_(iK)_}*). SNPs are encoded by allele count alleles (*X_(ig)_* ∈*{0, 1, 2}*). An ordinal logistic regression was considered to derive the probability than one SNP is associated to a multi-trait phenotype (Porter & O’Reilly, 2017). Permutation tests were performed to determine one adjusted significance threshold to detect false positives (Dudbridge & Gusnanto, 2008). A probability *P* < 5 × 10^-8^ was retained to consider one SNP as significantly implied in a multi-trait pigmentation association.

In single- and multi-trait GWAS, we controlled for population stratification by using nine distinct trout samples recognized by the length and number of ancestry tracts (Leitwein et al., 2018). Local ancestry tracts may integrate linkage disequilibrium patterns which are crucial in association studies (e.g. Shriner, 2017; Li, Kemppainen, Rastas, & Merilä, 2018). The nine groups include the two hatchery samples (Atlantic, Mediterranean), then seven distinct samples of wild-caught trout. Following Leitwein et al. (2018), wild-caught trout were grouped as follows: F1’s, F2’s, ‘early’ and ‘late’ backcrossed individuals, then samples of ‘pure’ wild individuals assigned to each of the three local populations. As some wild caught trout were identified as hatchery individuals (Leitwein et al., 2018), they have been grouped with individuals sampled in the hatchery type they have been previously assigned.

### 2.4 Mapping and annotation

CPLs detected by RDA and GWAS have been mapped on the Atlantic salmon genome (*Salmo salar*; Lien et al. 2016; Genbank assembly: GCA_000233375.4) and on the high density linkage map of *S. trutta* (Leitwein et al., 2017). We searched for genes located in a 25kb window upstream or downstream of each colour-associated marker (arbitrary range). When one association with a gene was detected, markers were assigned to coding or non– coding (intron, upstream or downstream sequences) gene regions, or other specific entities (transposable elements, pseudogenes). Using *S. salar* annotations as inputs, gene ontologies (GO) for biological processes and molecular functions were derived for genes associated to CPLs using QuickGO (https://www.ebi.ac.uk/QuickGO/). When annotations were unavailable on the salmon genome, a search was launched on UniProtKB (https://www.uniprot.org/uniprot/) for the protein sequence encoded by the gene. Only annotations with similarity >90% were retained.

Because many genes associated to melanocyte development are also implied in melanoma development when mutated (Patton, Mitchell, & Naim, 2010; Uong & Zon, 2010), we further performed a literature search in order to know if genes detected within the 25kb windows were previously mentioned in former studies. Queries were made using the gene ID (or aliases taken from GeneCards, https://www.genecards.org/) and a specific keyword. The retained keywords were: skin, melanocyte, melanophore, melanosome, melanoma (squamous, cutaneous and uveal), pigment/-ation, keratinocyte, chromatophore, iridiophore, follicle (hair or skin), nevus/-i, (epi)dermis, vitiligo, erythema/-tous, and sebocyte. This search was updated until the beginning of June, 2020, each time using the Web of Science, PubMed and Google Scholar.

### 2.5 Genomic differentiation

Distributions of *F*_ST_ values were established using all SNPs, then for the subset of CPLs found significant in the RDA and GWAS. Single-locus *F*_ST_ values were estimated considering five distinct trout samples: the two samples of hatchery fish, then trout caught in each of the three rivers within the Orb catchment.

### 2.6 – Comparison of individual assignment

In order to compare if individual assignment (probability of membership) ranked individuals similarly or not using all loci or CPLs only, we first established the probability of membership of individuals to the Atlantic population using independent CPLs defined in this study. Individual admixture proportions were computed with the LEA package (Frichot & François, 2015), using *K* = 2 (Atlantic and Mediterranean clades). We then used the probability of assignment established by Leitwein et al. (2018) that were based on individuals used in this study, using the same set of 40,519 RAD-loci, also considering *K* = 2. A Wilcoxon signed-ranked test between each assignment probability distribution was performed. Similar rankings of individuals in assignment will indicate that CPLs represent a proxy of genome-wide admixture, while a significant difference will be indicative of the reverse.

## 3. RESULTS

### 3.1 Redundancy analysis

Forward model selection showed that a model based on eight colour variables minimized the deviance in RDA *(Macrost, N.PR.Tot, Zeb, Fr.An, L.Lat, Fr.P, Diam.PN, N.PN.Tot*) (Table S3). Hereafter, results are reported for this model. Patterns of variation explained by the RDA were found to significantly structure the association between SNPs and pigmentation variables (*P* < 0.001). The first axis of RDA was found to represent 29.81% of the total inertia (Fig. 1), and was significant either (*P* < 0.001). RDA axis 2 was found to explain only a tiny fraction of observed variation for pigmentation variables (2.10%; Fig. 1), and be marginally significant (*P* = 0.051). As RDA is a constrained ordination method, constrained inertia (the percentage of variation explained by uncorrelated pigmentation variables) was found to represent 35.50% of the total inertia. The remaining portion is due to other factors. Axis 1 represented approx. 84% (ratio: 29.81/35.50) of constrained inertia, while axis 2 explained only approx. 6% (2.10/35.50) of inertia.

**Fig. 1:**
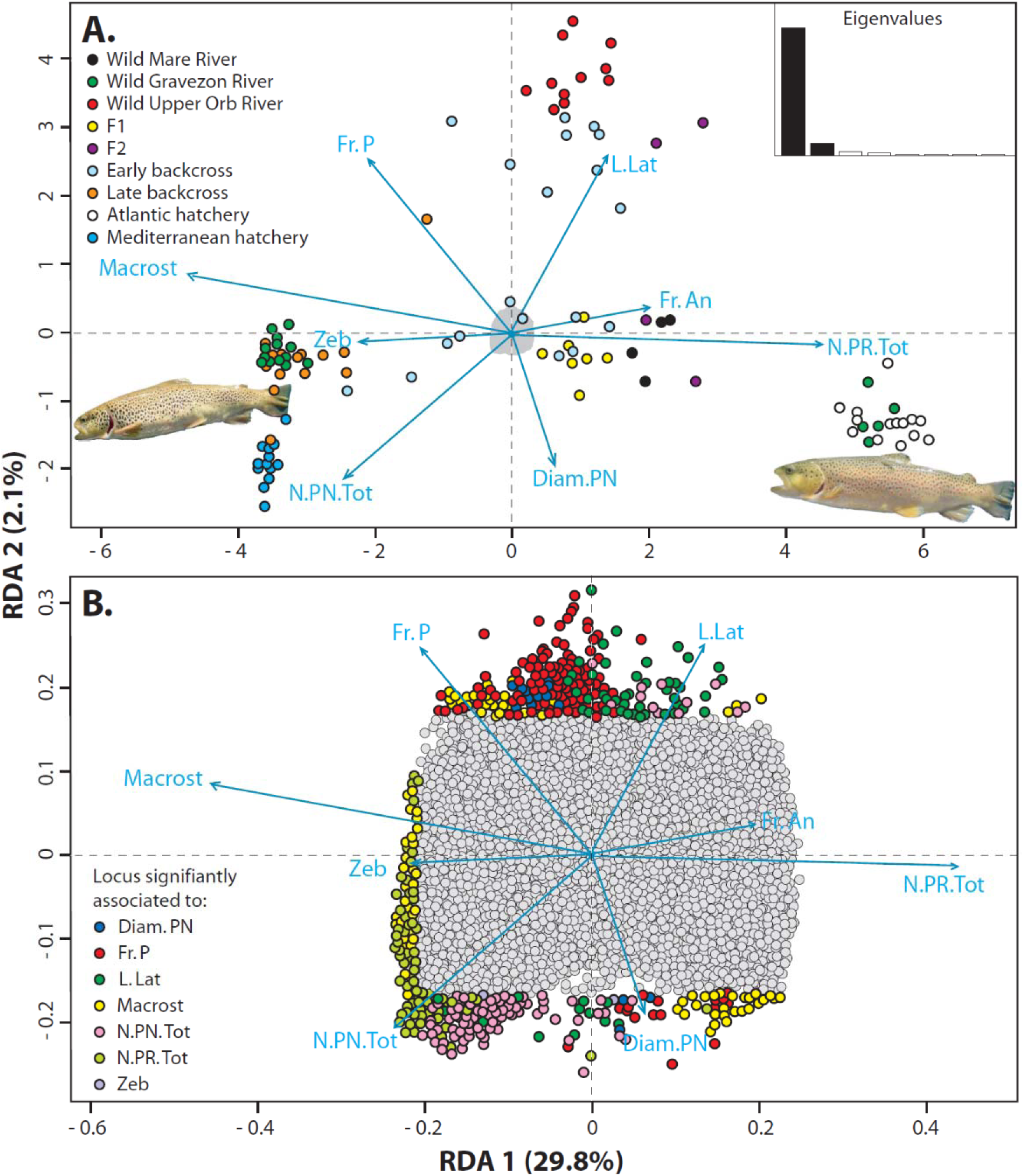
RDA triplots for significant or nearly significant canonical axes 1 and 2, respectively. Pigmentation-related variables retained by forward modelling (Blanchet et al., 2008) are represented by arrows; description of the retained model in Table S4. Length of arrows is proportional to the strength of correlation of each variable with individual axis. Arrows pointing to different direction indicate negatively correlated variables (*e.g. N.PR.Tot* and the large pre-opercular black stain *Macrost* for RDA axis 1). The percentage of variance associated to each axis is reported, as well as the eigenvalue graph for constrained axis. **(A):** Individual trout are positioned on the map with positioning of SNPs as a grey block to the center of the factor map. Individuals have been coloured according to the nine clusters defined by Leitwein, Gagnaire, Desmarais, Berrebi, & Guinand (2018) (top left); Clusters reflect the origin (hatchery *vs* wild) of trout, but also the degree of admixture of each of them as measured by co-ancestry tracts. **(B):** Zoom on the centre of the map to illustrate the position of SNPs. SNPs departing ± 2.5 standard deviates from the mean either on RDA axis 1 or RDA axis 2 have been coloured; a colour being associated to each pigmentation-related variable included in the model (insert). These SNPs are coloured with the variables they have been found the most significantly associated in the model. SNPs given as grey spots are within the ± 2.5 S.D. interval and not considered in this study. Percentages of variation explained by each RDA axis are reported. Fish are representatives of hatchery Mediterranean (left) and Atlantic (right) individuals used in this study.

RDA results are represented as a triplot in which individuals are positioned according to the relationship established between response and explanatory variables (Fig. 1). The ornamental trait *Macrost* (hereby a large pre-opercular black stain/spot) and the pigmentation trait *N.PR.Tot* (total number of red spots) explained 23.6% and 22.7%, respectively, of the total loading scores of phenotypic variables onto RDA axis 1. Results showed a clear distinction between colour patterns of Mediterranean and Atlantic hatchery fish (Fig. 1A). Atlantic hatchery fish were characterized by the total number of red spots, while the macrostigma spots, but also zebra marks (*Zeb*) and total number of black spots (*N.PN.Tot*) characterized Mediterranean hatchery fish along RDA axis 1. Results confirmed that few wild caught individuals from the Gravezon River were released Atlantic hatchery fish (Leitwein et al., 2018). Other individuals from the Gravezon River and ‘late backcrossed’ individuals have patterns more characteristic from hatchery Mediterranean fish (Fig. 1A; the hatchery was seeded by individuals caught in this river [see Materials and Methods section]). Other wild caught individuals are distributed between these extremes, with individuals identified as F1’s presenting intermediary positions between wild-caught and wild Mediterranean fish on one side, and Atlantic fish on the other side (Fig. 1A). The position of the wild caught Orb River individuals referred to the marginally significant second axis of the RDA and to specific variables (*L. Lat*, *Fr.P*) that were mostly unobserved in other fish (Fig. 1A). F2 individuals presenting these characters were effectively fished in the Upper Orb River, suggesting they possibly inherited colour characters expressed in this local population.

The ordination of SNPs by RDA is further detailed in Fig.1B. A total of 1,130 distinct loci (1.49% of 75,684 SNPs) were found significantly associated to the first two RDA axes (i.e. > 2.5 S.D. from the mean). Only 22 loci (0.03%) were significantly associated to each of the two axes for distinct phenotypic variables. Different numbers of SNPs were found associated with pigmentation variables: 299 with *Fr.P*, 269 with *N.PR.Tot*, 225 with *N.PN.Tot*, and 213 with *Macrost*. *Macrost* showed significance for loci associated to both first and second axis of RDA (yellow circles on Fig.1B). As axis 2 was found marginally significant and substantially affected by individuals from one single population, only SNPs associated to the first RDA axis will be considered further. This discarded loci associated to *Fr.P* that explained most of the variation for the second RDA axis. Three hundred and twenty SNPs were found associated to the main RDA axis, all of them being associated to *Macrost* and *N.PR.Tot*, these variables being negatively correlated (Fig. 1B).

As a complement to this analysis and because the threshold of ± 2.5 S.D. is somewhat arbitrary, we looked back at specific genes known to be involved in colour patterning in vertebrates and fish, and searched for their position within the distribution of loci over the RDA axis 1. We detected eight genes representing 30 SNPs for which sequencing reads were available (Fig. S2). One SNP from the *Mitf1* (microphthalmia-associated transcription factor 1) gene was close to the threshold considered in this study. The *Mitf1* gene is a master gene for melanocyte differentiation (e.g. Cheli, Ohanna, Ballotti, & Bertolotto, 2010) and, with a negative value on the first axis of the RDA, this SNP is associated to *Macrost*, thus the expression of the black colour.

### 3.2 Single and multi-trait GWAS

Only two pigmentation variables allowed for relevant LASSO model construction in single trait GWAS after selection of the penalized term using the cyclical descent procedure (Fig. S3): *N.PN.Tot* (total number of black spots: 17 candidate SNPs) and *N.PR.Tot* (total number of red spots: 9 candidate SNPs) (Table S4). These two variables were found significant in RDA, and negatively correlated (Fig. 1). Candidate SNPs from single GWAS models are reported in Fig. 2A.

**Figure 2:**
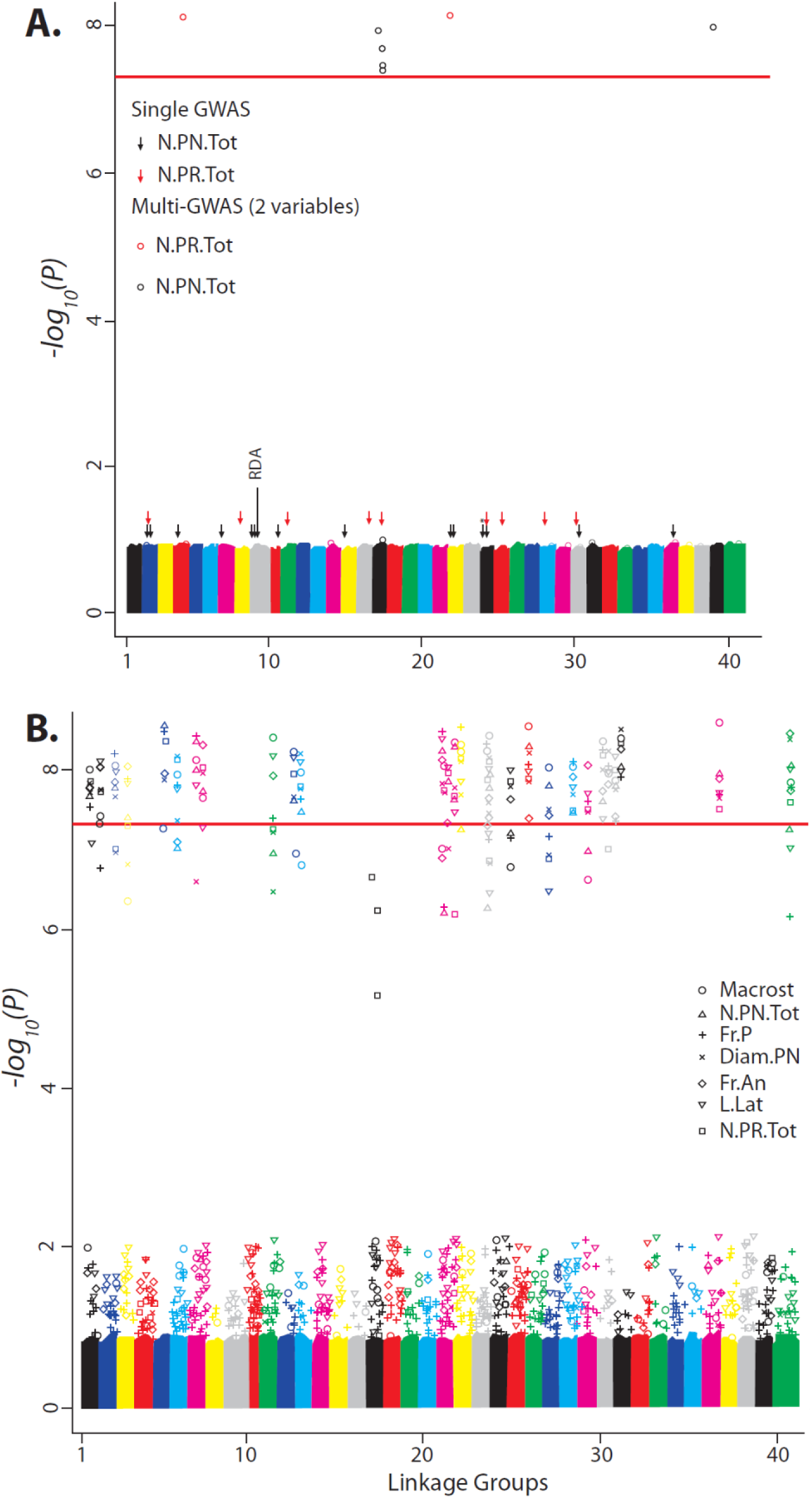
Manhattan plots of multi-trait GWAS models selected using the MultiPhen package (O’Reilly et al., 2012), plus results associated to the single-trait GWAS model used in this study. The abscissa axis represents the forty brown trout LGs (Leitwein et al., 2017), the ordinate axis reports values of *-log_10_(P)*, with *P* being the probability of one association between a variable and a SNP (adjusted significance threshold based on permutation, Dudbridge & Gusnanto, 2008). The red line indicates the 5 x 10^-8^ significance threshold retained in this study. **A:** Results for the multi-trait GWAS for the model using two variables: *N. PN.Tot* and *N.PR.Tot*. The seven SNPs found significant in this model are represented by circles. The results from single trait GWAS have been added to this panel to provide a full summary of results. In this case, approximate position of the RAD-loci associated to the *N. PN.Tot* (17 loci) and *N.PR.Tot* (9 loci) variables are indicated by black and red arrows, respectively. One asterisk indicates two loci to close of each other to be indicated by separate arrows. The locus associated to the *GJD2* gene on LG9 and also detected by the RDA is indicated by a longer arrow. **B:** Multi-trait GWAS based on the variables retained with the RDA except *Zeb* (31 candidate SNPs). The significance with each variable of the model is indicated by a different symbol. Each vertical line of symbols points to a single SNP. Some overlapping is possible. The full list of candidate SNPs detected using single and multi-trait GWAS are reported in Table S4. Variables labelled as in Table 1.

Two multi-trait GWAS models were retained (Fig. 2). As the single-trait GWAS, the first one jointly retained *N.PN.Tot* and *N.PR.Tot* as variables for which some SNPs were found associated. The second selected model considered seven of the eight variables formerly retained in the RDA (except *Zeb*). Seven and thirty-one SNPs were considered as significant in these multi-trait GWAS, respectively (Table S4).

Despite some similarities for the pigmentation and colour variables put forward by the different genotype-phenotype associations, only one single SNP (RAD-locus) was found in common between RDA and the single-trait GWAS for number of black spots (Fig. 2A). No other SNP was commonly detected by distinct genotype-phenotype association methods, and especially no SNP shared among GWAS models. It results in a total of 384 SNPs considered as CPLs putatively implied in pigmentation variation in brown trout (∼0.51% of the total number of SNPs; 320 coming from RDA and 64 from the single- and multi-trait GWAS models). These CPLs corresponds to 337 independent RAD-loci (0.83% of the total number of RAD-loci).

### 3.3 Genomic differentiation

The distributions of *F*_ST_ values for the 40,519 RAD-loci and the 320 CPLs associated to axis 1 of the RDA are reported in Fig. 3. The mean *F*_ST_ was estimated to *F*_ST_ = 0.286 [95% CI: 0.284, 0.287] for the full set of SNPs, while this estimate was *F*_ST_ = 0.575 [95% CI: 0.570-0.583] for the 320 CPLs detected with the RDA. Observed mean *F*_ST_ values were *F*_ST_ = 0.176 (min. 0.02 – max: 0.38) and *F*_ST_ = 0.025 (min.: 0.00; max. 0.20) for the single- and the multi-trait GWAS, respectively. Min/max values are reported rather than 95% CI because the low number of SNPs associated to phenotypic variables in each GWAS. As distributions do not overlap, the mean *F*_ST_ value of the 384 CPLs has no real meaning, but is *F*_ST_ = 0.547 [95% CI: 0.538 - 0.555].

**Figure 3:**
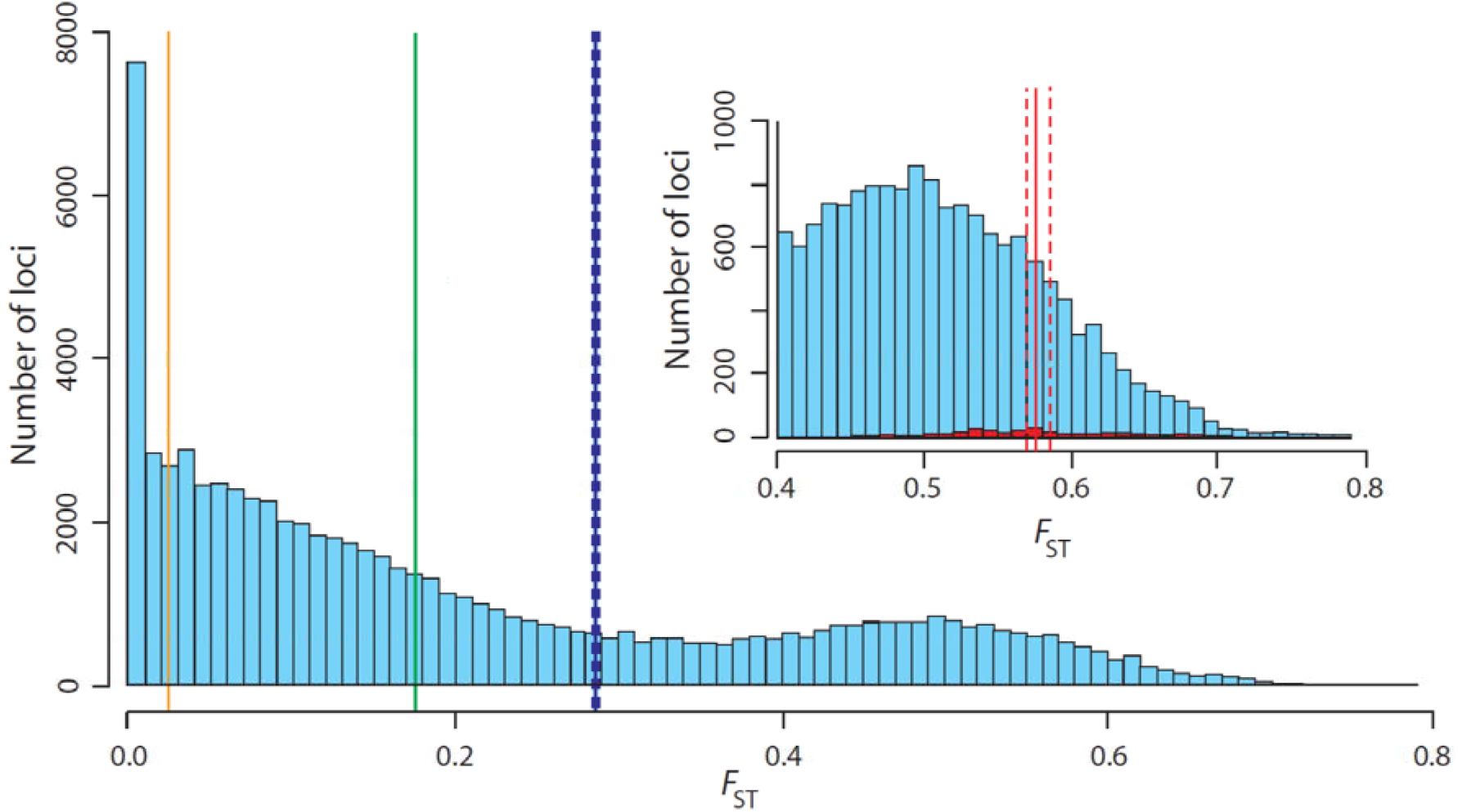
Distribution of *F_ST_* values for the 40,519 RAD-loci (blue) and for CPLs found significantly associated with RDA axis 1 (*N* = 320 loci, reported in red in the insert). For each data set, mean *F_ST_* values and confidence intervals are provided as full and hatched lines, respectively. Mean values for the single- and multi-trait GWAS approaches are illustrated by the green and the orange lines, respectively. No confidence intervals are reported for GWAS because of the low number of CPLs detected with these approaches. Details in the main text.

### 3.4 Mapping on *S. trutta* linkage groups and annotation

We considered only the 337 independent CPLs as linked SNPs provide redundant information. Three hundred independent CPLs were mapped onto 35 of the forty brown trout LGs defined in Leitwein et al. (2017) (Fig. 4), while 37 of them could not be adequately positioned on the high density linkage map of Leitwein et al. (2017). Their position relative to other RAD-loci suffered ambiguity. The distribution of these 300 CPLs on each LG varied from zero (LG13, LG14, LG15, LG20, LG39) to 86 (LG31). The 337 CPLs have been mapped on the Atlantic salmon genome and their annotations are given in Table S4. Among the 337 independent CPLs, ∼75.9% were found located in coding (43.4%) or so-called “regulatory regions” (32.5%; 17.8% upstream and 14.7% downstream genes) within the 25kb window retained around genes. This represents 245 loci associated to one gene (223 distinct genes; Table S4). Approximately 22.0% of the CPLs did not match any gene in the 25kb window, and ∼2% were found associated to pseudogenes or long non-coding (lnc) RNA (Fig. S4). LncRNAs might be important in fish colour patterning (Luo et al., 2019), but not considered further. Overall 89.86% (200 out of 223) of the genes in close vicinity of the CPLs detected in this study have been mentioned in the literature dealing with colour, or affecting integument patterning, differentiation or structure. Table S4 provides details on gene names, their position and the location of associated CPL; references mentioning each of these 200 CPLs are reported.

**Figure 4:**
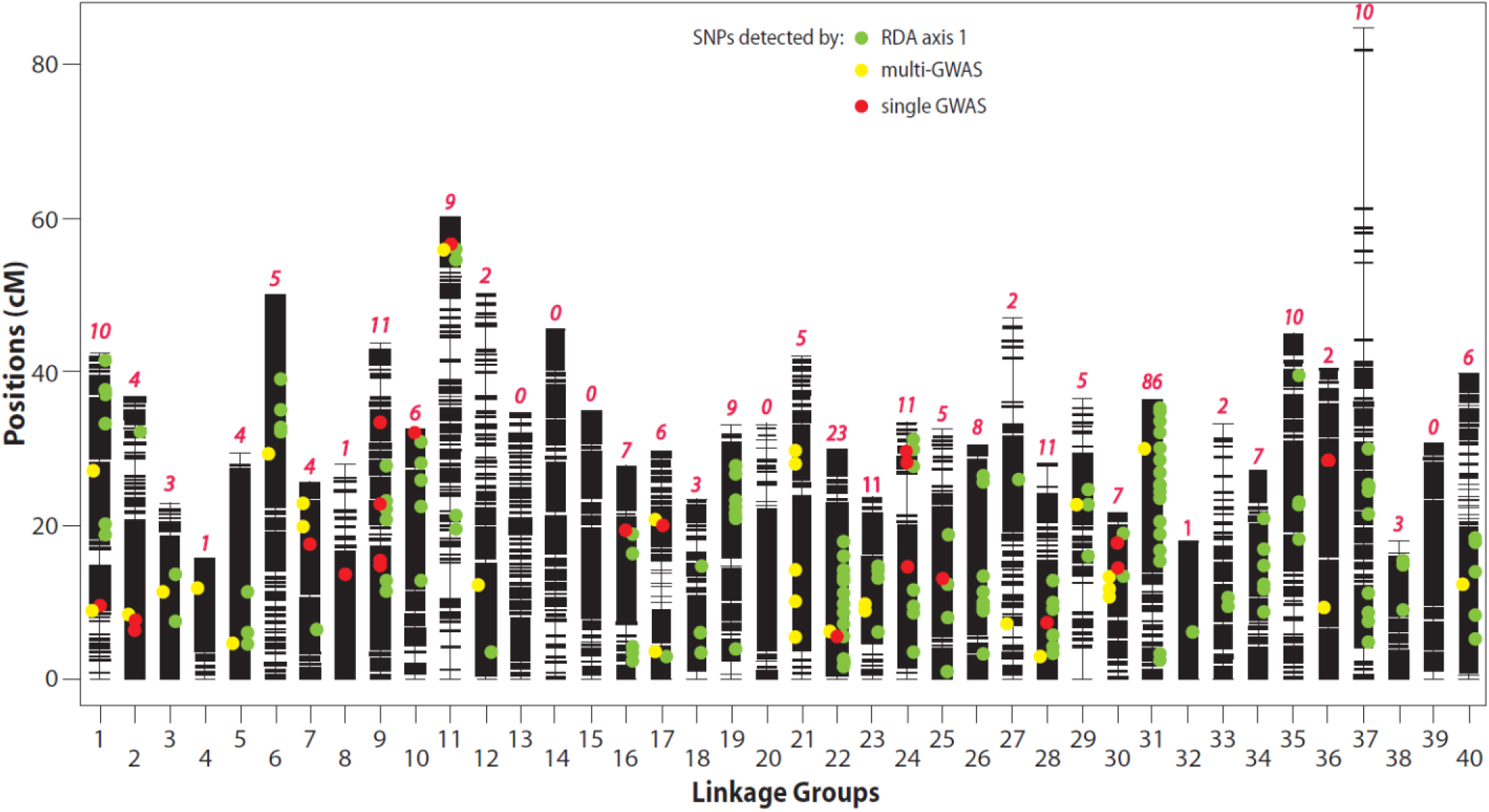
Positioning of the 300 mapped CPLs detected in this study on the forty LGs of the brown trout. Loci found associated at least one single phenotypic trait are indicted by green, red and yellow circles when detected with the RDA, single- or multi-GWAS analyses, respectively. Horizontal bars indicate the distribution of the full set of SNPs over LGs. The number of CPLs detected on each LG is indicated. As numerous loci are close to each other, circles may overlap.

An excerpt of twenty-four of these 200 genes (12.0%) is reported in Table 2 to summarize data. Among these genes, nine candidate genes were detected using GWAS (four by single-, and five by multi-trait GWAS) (Table 2). The RAD-locus detected by both RDA and single-trait GWAS for black spotting is located ∼21kb downstream of *GJD2* (Gap-junction protein Δ2 gene; located on LG9 of *S. trutta*; Table S4). This gene was formerly found differentially expressed in trout skin (Djurdjevič et al., 2019).

### 3.5 GO terms

GO terms for molecular functions and biological processes of the 337 CPLs are reported in Fig. S5. Calcium and metal ion binding were found to be the most representative molecular functions. GO-terms for biological functions highlighted cellular adhesion processes (Fig. S5A), signal transduction and translation, and transmembrane transport (Fig. S5B).

### 3.6 – Wilcoxon signed-ranked test on individual assignment

As expected because of large differences in mean *F*_ST_ values, individual rankings in probability of membership were found to significantly differ when using the whole RAD-loci established in Leitwein et al. (2018) and the independent CPL data sets defined in this study (Wilcoxon signed-rank test, *P* < 10^-5^). As CPLs were mainly associated to red and black spots (*N.PN.Tot*, *N.PR.Tot, Macrost*) in the GWAS and the RDA, this suggests that such patterns are not reliable indicators of genome-wide admixture, while frequently, but intuitively used by managers.

## 4. DISCUSSION

Because of the importance of visual information in natural and sexual selection, we aimed to investigate the genomic basis of body colour patterning between trout from the Mediterranean and Atlantic clades recognized in this species. The second one is heavily used for stocking practices and has been introduced worldwide, including Mediterranean watersheds (Elliott, 1989; Budy et al., 2013). Based on findings by Leitwein et al. (2018), admixture (ancestry) patterns were included in our study and improved the detection of significant associations as shown in previous studies (e.g. Shriver et al., 2003; Daya et al., 2014; Pallares, Harr, Turner, & Tautz, 2014; Brelsford, Toews, & Irwin, 2017), including fish (Malek et al., 2012). This information allowed for the detection of CPLs associated to candidate genes formerly identified in pigmentation studies. These CPLs complement genome-wide gene expression studies by Djurdjevič et al (2019) and improve knowledge on the genomic basis of pigmentation in the brown trout. While pigmentation patterns are important to evaluate the impact of hatchery individuals into wild trout populations (Mezzera et al., 1997; Aparicio et al., 2005), it is premature to rely on phenotypic colour patterns as a proxy of genome-wide admixture to promote management decisions.

### 4.1 Associations promote spotting traits…

Genotype-phenotype associations used in this study principally put forward the numbers of red and black spots and the presence/absence of the black pre-opercular spot as the main pigmentation variables that aimed to distinguish Mediterranean and Atlantic trout. This was expected as these variables have been repeatedly shown to characterize pigmentation differences between Atlantic or Mediterranean trout in studies previously conducted in France (Poteaux & Berrebi, 1997) and Spain (Aparicio et al., 2005). These traits were not always found in Italian populations, and others may prevail locally (e.g. patterns of the adipose fin) (see Lorenzoni et al., 2019). While focusing on few local French populations, results presented in this study thus reflect most of the former results obtained for these two clades at a broader scale. Nevertheless, as for Italian populations, model selection showed that potentially more variables might be considered at the local scale. Spotting traits do not uncover the full panel of pigmentation variation present in a polytypic species like brown trout. Further works participating to establish the genotype-phenotype association for colour pattern in brown trout should integrate other tools and variables than used in this study. Recent software developments may participate to improve this issue (Endler, Cole, & Kranz, 2018; Van Bellenghem et al., 2018; Chan, Stevens, & Todd, 2019), including additional data on hue, reflectance or brightness.

### 4.2 … but distinct loci involved in pigmentation variation

If the different genotype-phenotype association methods used in our study retrieved the same pigmentation variables, CPLs detected by RDA and GWAS appeared clearly distinct with exception of *GJD2* formerly studied in trout (Djurdjevič et al., 2019). Indeed, the identity of loci considered associated to a single and/or multi-trait phenotype, their number, and their respective levels of genomic differentiation did not overlap. This is expected as models behind methods are different and may direct the analyses towards specific findings. For example, we already mentioned that the shrinkage procedure in LASSO aimed to keep only SNPs with large effects while discarding others, also explaining why far less CPLs come from GWAS. It is also widely known that results from single-trait GWAS lack power when few individuals or SNPs are considered (e.g. Schielzeth & Husby, 2014), especially when considering wild populations (Kardos, Husby, McFarlane, Qvanström, & Ellegren, 2018; Santure & Garant, 2018) and polygenic variation (Berg & Coop, 2014). These drawbacks were moderated by consideration of population stratification based on ancestry patterns. In the opposite, GWAS retrieved loci with low and sometimes null *F_ST_* values that RDA ignored while balancing selection is crucial to explain pigmentation variation (e.g. Croucher, Oxford, Lam, & Gillespie, 2011; Lindtke et al., 2017; Schweizer et al., 2018). We do not expand further. A sub-section below is devoted to RDA that, despite caveats, remains potentially neglected in association studies.

We prefer to adopt a pragmatic view: RDA and GWAS detected a large proportion of markers formerly mentioned in the literature on skin and pigmentation. Over the 337 independent CPLs retained overall, 245 loci mapped within a 25kb window downstream or upstream 223 distinct genes. Two hundreds of them were formerly mentioned in our literature survey regarding issues and processes dealing with colour patterning or skin-related issues. The GO terms highlighted by our study for both molecular functions and biological processes are crucial to pigmentation patterning. Calcium binding is known to be important for melanogenesis (Bush & Simon, 2007; Bellono & Oancea, 2014; Jia et al., 2020). For example, the *SLC45A2* gene detected in this study that encodes for a transporter protein involved in the production and regulation of melanin is regulated by calcium binding levels (Ginger et al., 2008; Bellono & Oancea, 2014). Melanin is also known to deeply interact with metal ions at various steps of melanogenesis (Hong & Simon, 2007). Metal ion binding is necessary in the integrin-mediated signaling pathways whose importance in colour patterning is well-known (Kelsh et al., 2009; Klotz et al., 2018). Zinc binding is critical to control melanocyte migration to the epidermis (Denecker et al., 2014) and microtubule activity for melanosome trafficking within melanocytes (Aspengren, Hedberg, Sköld, & Wallin, 2009). Other functions such ATP- or DNA-binding activities participate to melanosome/melanocyte biology and regulation (Levy et al., 2006; Heimerl, Bosserhof, Langmann, Ecker, & Schmitz, 2007). DNA-binding especially concerns transcription factors that promote growth, survival and differentiation of melanocytes. The main biological functions highlighted in this study (cellular adhesion processes, signal transduction and translation, transmembrane transport) are involved in chromatophore or melanocyte development, interactions among chromatophores or between melanocytes and other cell types (fibroblasts, keratinocytes) (Yamaguchi, Brenner, & Hearing, 2007; Raposo & Marks, 2007; Kelsh et al., 2009; Mahalwar, Singh, Fadeev, Nüsslein-Vollard, & Irion, 2016; Nüsslein-Völlard & Singh, 2017). Cell adhesion processes are illustrated by the numerous calcium-dependent (proto)cadherin genes found in this study (*CDH4*, *FAT1*, and *PCDH* genes), as well as integrins (*ITGAV*) (Tables 2 and S4). Protocadherins (e.g. *PCDH10A*, *PCDHAC2*) are essential to fish skin patterning (e.g. Williams, Hsu, Rossi, & Artinger, 2018; Du et al., 2019).

Each association method reported classical candidate genes associated to pigmentation. The RDA reported the above-mentioned (proto)cadherins, but also the *ZIC4* or *SLC7A11* genes. *ZIC4* participates to the dorso-ventral patterning of medaka (Ohtsuka et al., 2004). The observed trait association with *SLC7A11* – a gene controlling pheomelanin production in mammals (Chintala et al., 2005) - is another report in fish after carp (*Cyprinus carpio*; Xu et al., 2014; Jiang et al., 2014) and the ‘red’ tilapia (Zhu et al., 2016; Wang et al., 2019). GWAS detected some classical players of pigmentation and body colouration in vertebrates (e.g. *SOX10, PEML, SLC45A2*), including fish (Schonthaler et al., 2005; Hou, Arnheiter, & Pavan, 2006; Kelsh et al., 2009; Singh & Nüsslein-Vollard, 2015). Indeed, *SOX10* influences colour patterning during zebrafish development (Elworthy, Lister, Carney, Raible, & Kelsh, 2003; Greenhill, Rocco, Vibert, Nikaido, & Kelsh, 2011; Nüsslein-Vollard & Singh, 2017). *SLC45A2* is involved in proton-transport and osmoregulation of melanosomes resulting in colour dilution as observed in medaka (Fukamachi, Shimada, & Shima, 2001) and other organisms (e.g. Xu et al., 2013; Adhikari et al., 2019; Sevane, Sanz, & Dunner, 2019). *PEML* (premelanosome protein) contributes to eumelanin deposition and colour dilution in tetrapods (Raposo & Marks, 2007; Ishishita et al., 2018), and found to be involved in colouration of the koi carp (Luo et al., 2019) and a cichlid (Ahi & Sfec, 2017).

Finally, few CPLs are associated to genes involved in background colour adaptation (*GABRA2*, gamma-aminobutyric acid receptor subunit alpha-2-like; Bertolesi, Vazhappill, Hehr, & McFarlane, 2016), or light-induced colour change (*MTRR*, methionine synthase reductase-like; Jones et al., 2018; *FGFR1*, fibroblast growth factor receptor-like 1; Czyz, 2019). *FGF* signaling is essential to the developmental patterning of skin and scales in fish (Aman, Fulbright, & Parichy, 2018), but also eyespots of butterfly wings (Özsu & Monteiro, 2017). This raises interesting questions regarding the conservation of mechanisms involved in spot formation in different developmental contexts.

Overall, results established for the brown trout identified both key genes and an array of far less-known genes involved in biological and molecular processes related to the regulation of colour and skin pigmentation patterning.

### 4.3 A highly polygenic architecture with a role for drift

With 337 independent CPLs distributed over 35 LGs and a fraction of them with well-established relationships in modulating colour and pigmentation patterns, a polygenic architecture of body colour patterning is satisfied in the case of brown trout. Informative CPLs participating to genomic architecture of pigmentation in trout are shared in both coding and regulatory regions. This partitioning is now often reported in the literature (e.g. She & Jarosz, 2018), but still remains relatively poorly investigated in fish (Peichel & Marques, 2017, but see Jones et al., 2012). Within pigmentation studies, literature emphasizes both the relative roles of coding (Protas & Patel, 2008; Uy et al., 2016) and regulatory variation to limit pleiotropic costs during colour pattern establishment and evolution (Larter, Dunbar-Wallis, Berardi, & Smith, 2018; Toomey et al., 2018). More coding changes than loci dispersed in regions that might be the basis of regulatory changes were detected in trout, but, due to the reduced representation of the genome provided by ddRAD sequencing, it is difficult to appreciate the relevance of this observation. It might be simply due to the choice of restriction enzymes that returned a peculiar genomic distribution of CPLs. Furthermore, we provided only a loose definition of upstream or downstream regulatory regions (a 25kb window) that, undoubtedly, has to be refined in further investigations to better decipher, e.g., *cis*-regulatory processes.

This polygenic architecture contrasts with former studies that supported oligogenic models or showed the likely presence of few major QTLs in trout and salmonids (Blanc et al., 1982, 1994; Boulding et al., 2008; Colihueque, 2010), but also in other fish species (*n* < 8 QTLs; Borowsky & Wilkens, 2002; Miller et al., 2007; Magalhaes & Seehausen, 2010; Greenwood et al., 2011; Malek et al., 2012; O’Quin et al., 2013; Yong et al., 2015). However, few QTLs are considered not sufficient to explain all causal variation for pigmentation differences (Greenwood et al., 2011) and comparison with our study should be carefully considered. First, we did not use fish from controlled crossing experiments and our ability to detect any QTL is null. Second, we did not focus on a given body part (e.g. fin) or a pigmentation character (stripe, bars, belly spots) as done in other studies (e.g. Ahi & Sfec, 2017; Roberts et al., 2017), including trout (red spots in Blanc et al., 1994). We explored the complexity in trout pigmentation in order to detect the most important body pattern-related variables relevant to our samples. This automatically increased the panel of loci involved in body colour pigmentation, and, as in this study, when more pigmentation traits were considered, the number of pigmentation-related QTLs was found far higher and distributed over more LGs than when studying only one or few traits (*n* > 20; Tripathi et al., 2009; Albertson et al., 2014).

One increased genomic basis to pigmentation patterns was also reported when drift was present (Protas, Conrad, Gross, Tabin, & Borowsky, 2007), and drift is common in trout because of small local effective population sizes (*N_e_*) (e.g. Palm, Laikre, Jorde, & Ryman, 2003; Charlier, Laikre, & Ryman, 2012). This relies on the classical contrast between the quantitative trait nucleotide (QTNs) and the infinitesimal models of trait variation (e.g. Rockman, 2012). When stochastic processes are of prime importance, small infinitesimal segregate in many genomic regions and might be recruited in the mechanisms and programmes of leading to trait variation (Rockman, 2012). This may translate to a genomically diffuse information to specific phenotypes at the local population level as observed in trout. This is illustrated by loci associated to RDA axis 2 that are population specific (Upper Orb River) and concern peculiar trait associations. The highly polytypic brown trout may illustrate one intermediate situation, in which some master genes are detected at the clade level by both GWAS and RDA and are associated to recognised spotting traits, while ‘small players’ mostly detected by RDA illustrate patterns specific to local populations, more locally expressed by other traits *(L. Lat*, *Fr.P*). If the big picture provided by master genes should be investigated further, ‘small players’ cannot be ignored.

### 4.4 RDA: A neglected method to screen for genotype-phenotype associations

Our findings illustrate the ability to investigate genotype-phenotype associations with a constrained ordination method like RDA associated to accurate modeling of phenotypic variables. Jombart, Pontier and Dufour (2009) already mentioned that RDA was neglected in association studies despite now recognized desirable properties to limit false positives (Capblanc, Luu, Blum, & Bazin, 2018; Forester et al., 2018) and robustness to recombination rate variation (Lotterhos, 2019). Effectively, the use of RDA to investigate genotype-phenotype associations remains seldom (Talbot et al., 2017; Vangestel, Eckert, Wegrzyn, St. Clair, & Neale, 2018; Carvalho et al., 2020), while it became a standard in genotype-environment association studies (Forester, Lasky, Wagner, & Urban, 2018), including trout (Bekkevold et al., 2019). Basically, RDA allowed for distinguishing the part of variation collectively explained by RAD-loci and independent pigmentation variables (∼35% of observed trait variation). A large portion of observed pigmentation might be noise or be environment-dependent. Studies in salmon (Jørgensen et al. 2018) and trout (Westley et al., 2013) showed that plasticity may indeed prevail in explaining pigmentation patterns. While it discards markers under balancing selection, a beneficial feature of RDA is to possibly focus on SNPs that may explain for causal variation with at least one phenotypic variable, and not only using outlier loci with the highest *F*_ST_ values, but without demonstrated association to a phenotype. For example, in a study dealing with pigmentation variation, Neethiraj, Hornett, Hill, and Wheat (2017) used genome scans tools to identify outlier loci, then *a posteriori* searched for association of these loci with pigmentation. If RDA remains affected by some false positives as are genome scans, it certainly targets more relevant loci as illustrated by the high number of CPLs retrieved from the literature in this study. Gautier (2015) proposed a model based on Bayesian outlier detection model that may use a color variable as a covariate, but it is limited so far to a single covariate when model-based selection prior to RDA might allow for considering several ones concurrently. Adding on Jombart et al. (2009), RDA or related methods (e.g canonical correlation analysis, Porter & O’Reilly, 2017) deserve more attention in association studies.

### 4.5 Implications for management

Contrary to birds (Hanna et al., 2018; Billerman et al., 2019) or mammals (e.g. Anderson et al., 2009; Fulgione et al., 2016), how differences in body pigmentation are related to differentiation and admixture was poorly studied in fish and was not related to conservation and management issues (Malek et al., 2012; Meier et al., 2018, but see Boulding et al., 2008). Using mitochondrial and allozymic markers, Aparicio et al. (2005) paved the way to use pigmentation as a proxy of admixture between Atlantic and Mediterranean trout and to address such issues. Indeed, body pigmentation patterns are often important to anglers and managers to characterize local trout populations and their “integrity”. Earlier studies addressing this issue using old-generation markers indicated that pigmentation might represent such a proxy (Mezzera et al., 1997; Aparicio et al., 2005; see Delling et al., 2000 for *S. t. marmoratus*), but this issue was not addressed with SNPs. In this study, RDA showed that ‘late backcrossed’ individuals clustered very close from the local Mediterranean hatchery strain and wild individuals of the Gravezon River that were used to seed this strain. This suggests that the pigmentation pattern of these latter individuals came back close to the local Mediterranean patterns and that counter selection of Atlantic pigmentation alleles or other changes (e.g. Endler, Betancourt, Nolte, & Schlötterer, 2016) occurred. However, different rankings in assignment between this study for CPLs and the full RAD-loci data set (Leitwein et al., 2018) also suggest that this set of CPLs is not a proxy of genome-wide admixture in this catchment. Managers could build a policy based on wild-caught individuals that exhibit the ‘local pigmentation’ whereas trout remain largely admixed with foreign alleles. If this mismatch between rankings is due to neutral alleles, admixture would hopefully not constrain the adaptive potential of populations, and pigmentation-based management policies that limit the genotyping costs be sufficient. On the contrary, if a mismatch occurs and that CPLs do not mirror admixture at locally-adapted fitness effect loci, using pigmentation as a proxy of local genetic integrity alone might be at risk despite apparent counter-selection. Distinguishing between these situations is a difficult task, but, at that time, heavy reliance on the ‘*local is best*’ paradigm (i.e. considering that individuals presenting the ‘local’ phenotype are more adapted to local conditions) might be harmful for population fitness (Broadhurst et al., 2008; Kronenberger et al., 2018). We are aware that the list of CPLs we propose is a list of candidate markers that has to be refine for further testing against genome-wide admixture, but, in the meantime, managers should adopt a precautionary approach when evaluating population ‘integrity’ based on body pigmentation.

### 4.6 Limits and future directions

While providing strong results, comparisons using individuals from more natural populations, clades and/or subspecies within the *S. trutta* complex remain necessary in order to better investigate and elucidate the evolutionary history of trout regarding pigmentation patterns and to refine the CPL list. DdRADseq used in this study provides a reduced genome representation targeting only a tiny portion of the genome that is considered either to successfully target (Belsford et al., 2017), or miss important loci responsible for pigmentation patterns (Gauthier et al., 2020). Thus, further RNA sequencing or gene expression studies, together with more standard association studies in control settings that, as for other traits (e.g. Sinclair-Waters et al., 2020), may provide estimates of QTL number, variation in locus effect sizes and more accurately pointing key master genes seem necessary to improve findings. However, the particular settings of many natural trout populations (low *N_e_*, high genetic drift and reduced gene flow) have to be taken into account to explain pigmentation diversity observed in the wild. Pigmentation issues represent one outstanding case to investigate genotype-phenotype-environment relationships, and increasing knowledge on such relationships is crucial in salmonid conservation (Waples et al., 2020). As thousands of trout pictures are taken each year and tissue samples might be easily collected, the development of conservation phenogenomics programmes involving scientists and practitioners should be motivated.

## Supporting information

Table Suppl. Mat. S4 that cannot be accurately with other supplementary materials included

## Acknowledgements

This work benefited from the Montpellier Bioinformatics Biodiversity (MBB) platform from the LabEx CeMEB, an ANR “Investissements d’Avenir” program (ANR-10-LABX-04-01). J. Pouzadoux and A. Jourdan are acknowledged for inputs. ML was partly supported by a grant from LabEx CeMEB.

## Data accessibility

Genomic data are from Leitwein et al. (2018) and available at NCBI Short Read Archive under the study accession SRP136716. The final VCF and the PLINK files for *S. trutta* LGs are available under the study accessions SRZ187687 and SRZ187688. Upon acceptance, trout pictures and phenotypic data will be deposited on Dryad or another repository.

## Authors’ contribution

TV, ML and BG analyzed data and drafted the manuscript. JML acquired phenotypic data. ED provided support to molecular data analysis. ML, PB and BG designed the study. All authors revised drafts and approved the final version.

## Notes

### Competing Interest Statement

The authors have declared no competing interest.

