## Supplementary material for "Spotting genome-wide pigmentation variation in a brown trout admixture context": Captions and Supplementary Materials

**a brown trout admixture context**

**______________________________________**

**Captions of Supplementary Tables**

**Page 4:	Table S1 : Phenotypic variables used in this study (*N* = 30).** The nature and the coding of each quantitative and semi-quantitative variable are reported.

**Page 5:	Table S2 :** **Pair-wise correlation between phenotypic variables**. Following Blanchet et al. (2008), only uncorrelated variables (green cells) have been conserved for further analysis and model selection, leading to *N* = 11 variables left out of the 30 initially considered (Table S1).

**Page 6:	Table S3 : Best RDA model based on forward selection following Blanchet et al. (2008).** The best fitted model explaining SNP variation depends on a model that includes 8 phenotypic variables for color patterns (variable labels in Table S1). This model minimizes the Aikake Information Criterion (AIC). The table reports the effect of adding (**+**) or deleting (**-**) some variables to this model. In each case, AIC is increased indicating that some variables were not useful to the model (*Fr.D, Parr, Oc.PN*). Deletion of variables included in the model also increased AIC proving they were useful to the model as assessed by the associated probability. *Df* : degree of freedom ; *F* : F-statistic, *Pr(>F)* : associated probability.

**Separate file:	Table S4 : Gene names, gene symbols associated the nearest gene of each color pattern-associated loci (CPALs)** on the LGs the brown trout (*Salmo trutta*) linkage groups (LG) defined by Leitwein et al. (2017; *G3* **7:** 1365-1376. doi:10.1534/g3.116.038497). The correspondence with the Atlantic salmon (*S. salar*) genome is reported (Lien et al. 2016; *Nature* **533:** 200-205. doi: 10.1038/nature17164). For each locus, the association method that allowed for its detection is reported in the first column, the nature of the loci (coding, upstream or downstream the closest gene within a 25kb window) is reported. Blank cells indicate no match. GO terms are provided and a second match with another gene also included in the 25kb window considered in this study is reported when it has been detected. Doi or PMID identifiers of studies that reported the identified gene as having a role in colour patterning or a suite of key words dealing with skin (see second sheet of the table for the full list) are reported in green columns on the right side of the table. Few studies (e.g. Ph.D. thesis) without doi might also be reported. Orange cells refers to genes listed in Table 1 of the main text and represent the genes found in this study that are the most commonly reported in the literature dealing with color and color patterning in animals. The blue line points to the *GJD2* gene known to be differentially expressed in trout skin (Djurdjevič et al. 2019; *BMC Genomics* 20: 359. doi:10.1186/s12864-019-5714-1) and detected by two independent association method in this study. The last line of the table summarizes findings.

**______________________________________**

**Captions of Supplementary Figures**

**Page 7:	Figure S1: Illustration of the phenotypic variables used in this study.** Each variable was acquired separately on the left flank of trout.

**Page 8:	Figure S2:** Loading scores along axis 1 of the RDA (*y*-axis) of SNPs (*N* = 30) located in 8 pigmentation- or colour-related genes taken from the literature and for which sequencing reads were available. The order of genes is arbitrary and each gene is represented by a single color. The red lines represent the ± 2.5 stand deviation threshold considered in this study. None of these loci is close to this threshold except one SNP associated to the *Mitf1* gene, a master gene for melanocyte differentiation and expression of the black colour. Symbols of candidate genes with sequencing reads are reported. Support for their role in colour patterning can be found in, e.g., Raposo and Marks (2007; *Nat. Rev. Mol. Cell Biol*. **8:** 786-797. doi: 10.1038/nrm2258), Kelsh et al. (2009; *Semin. Cell Dev. Biol*. **20:** 90-104. doi: 10.1016/j.semcdb.2008.10.001) or Kronforst et al. (2012; *Pigment Cell Melanoma Res*. **25:** 411-433. doi: 10.1111/j.1755-148X.2012.01014.x) for *Dct*, *Mitf*, *Kit*, *Tyrp1*, *Pomca1*, *Pomca2*, *Pomcb,* while *Scg2a* was especially found differentially expressed in the skin of the marble trout *S. trutta marmorata* (e.g. Sivka *et al.* 2013; *Comp. Biochem. Physiol. D Genom. Prot.* **8:** 244-249. doi: 10.1016/j.cbd.2013.06.003). Note that the pattern for *Mitf1* is similar to variation of *F_ST_* along a chromosome: 10 SNPs (8 RAD-loci) were present in *Mitf1* with only one close to the significance threshold. Loading scores along RDA axis are proportional to *F_ST_* values.

**Page 9:	Figure S3:** Illustration of the estimation of the penalty parameter *λ* by the cyclical descent procedure proposed by Friedman et al. (2010; *J. Stat. Softw*. **33:** 1-22. PMID: 20808728) for use in the LASSO model used and single-GWAS analysis. The value of log(*λ*) minimized the mean quadratic error and the number of *β_j_* coefficients used in the model (the number of *β_j_* > 0 coefficients is reported at the top of the figure).

**Page 10:	Figure S4**: Distribution of CPALs found in this study in coding gene regions, or upstream/downstream genes (a 25kb window was considered). Positions are based on the mapping of sequencing reads onto the annotated Atlantic salmon genomes. In few cases (~2%), CPALS were found associated to long non-coding (lnc) RNA or pseudogenes, and for ~24% of them, intergenic (i.e. outside the 25kb window considered in this study to be associated to a gene).

**Page 11:	Figure S5:** Main GO-terms associated to the CPALs defined in this study. Results are reported for **(A)** molecular functions, and **(B)** biological processes.

**Table S1**

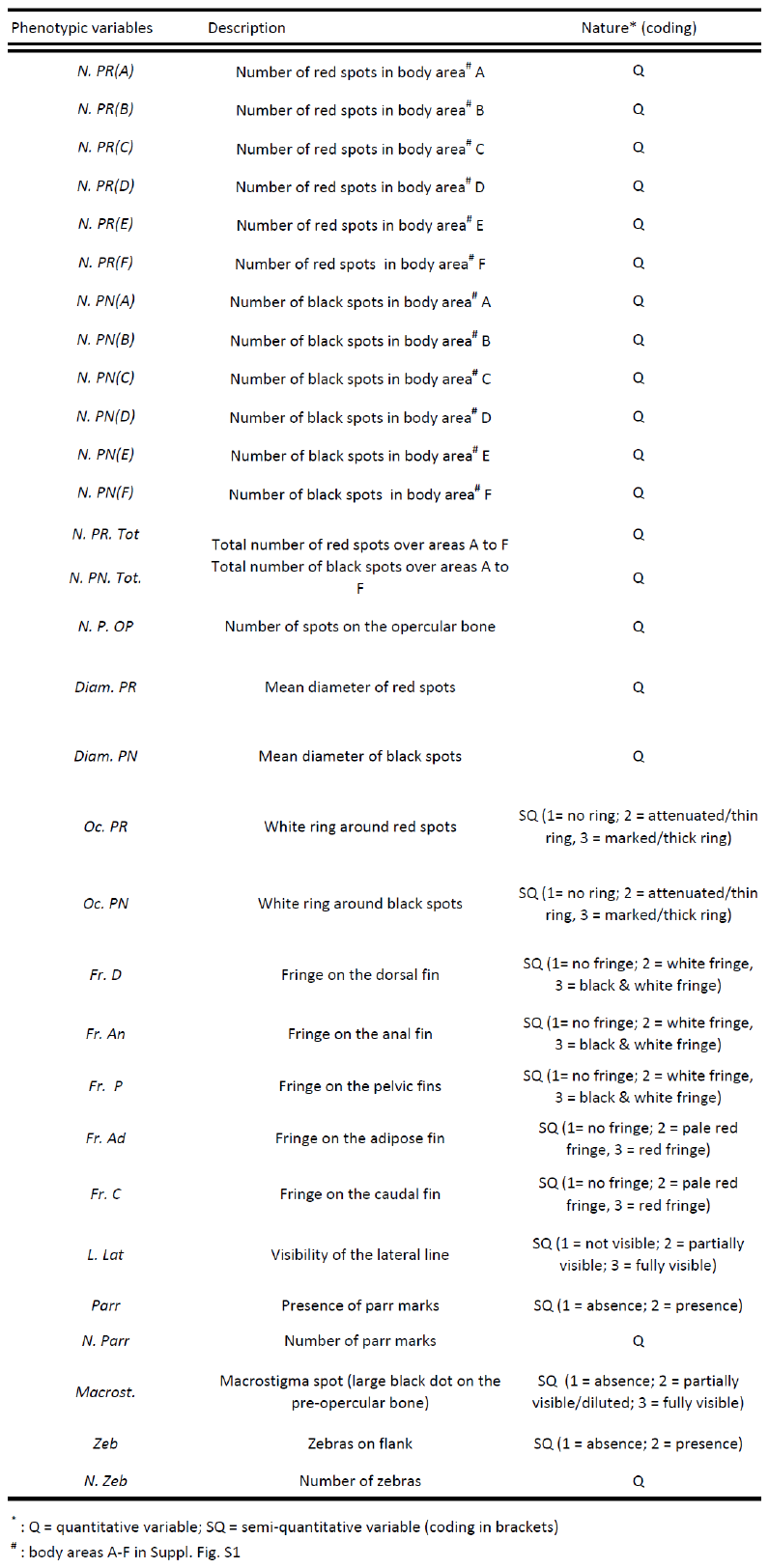

**Table S2:**

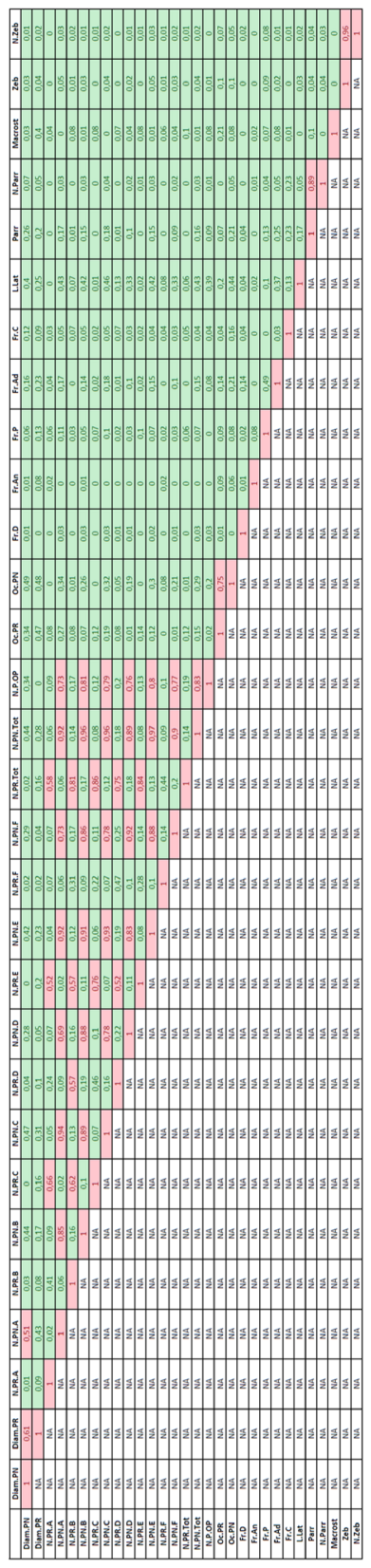

**Table S3**

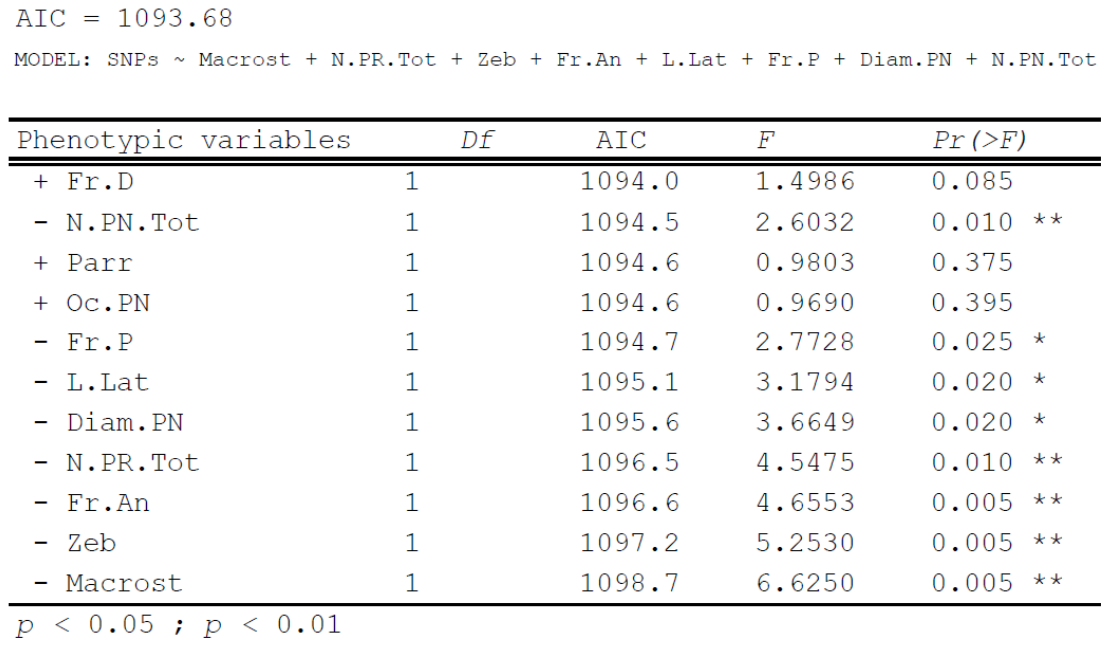

**Table S4:**

Too large for this material. Provided as a separate Excel file or provided upon request to:

_________________

**Figure S1:**

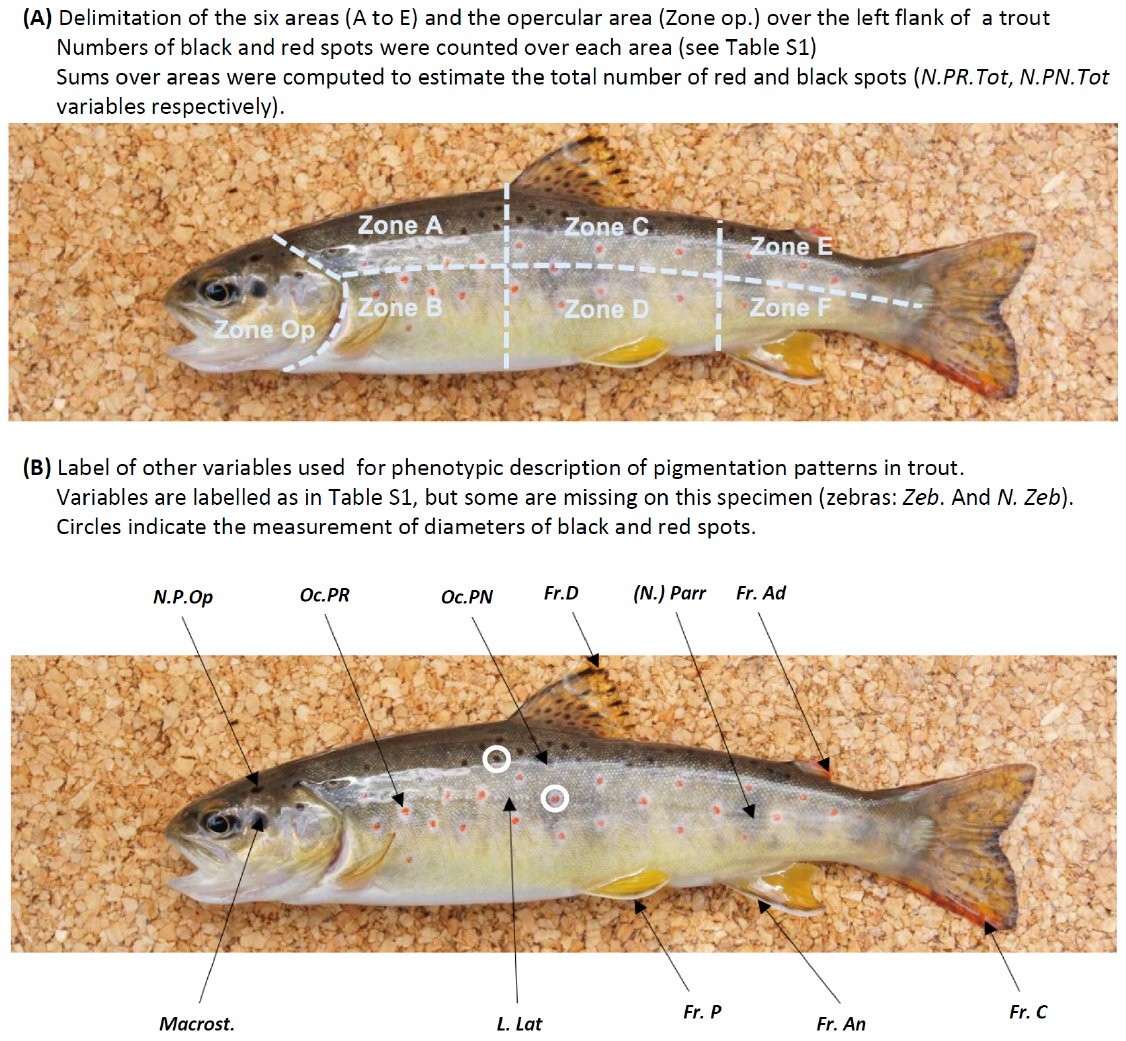

**Figure S2:**

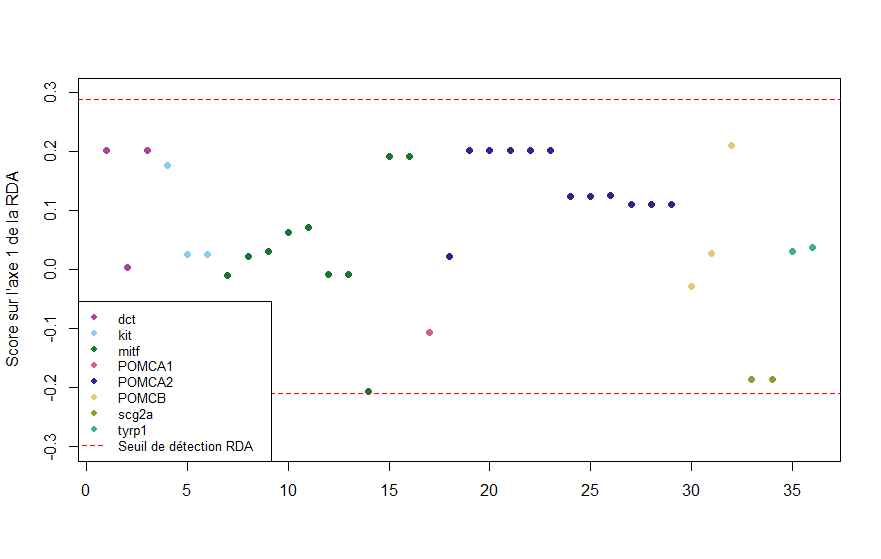

**Figure S3 :**

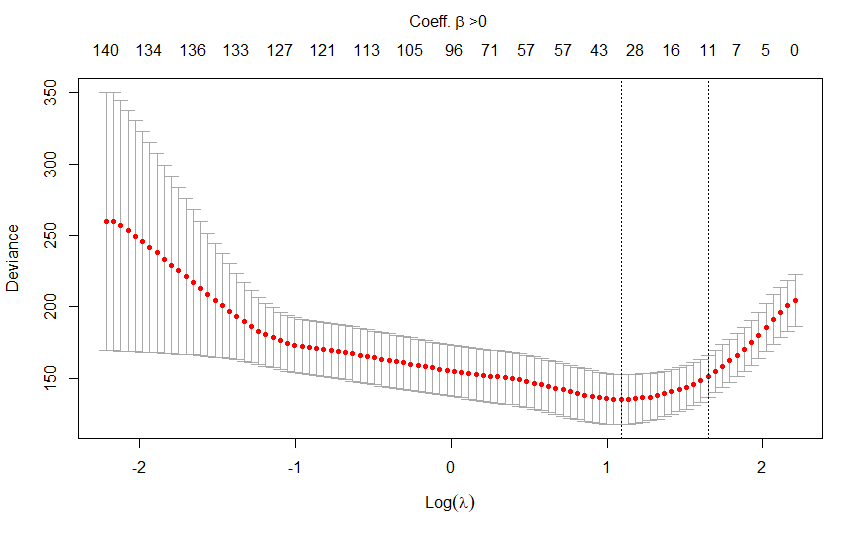

**Figure S4:**

Coding

Outside the

25kb window

retained in

this study

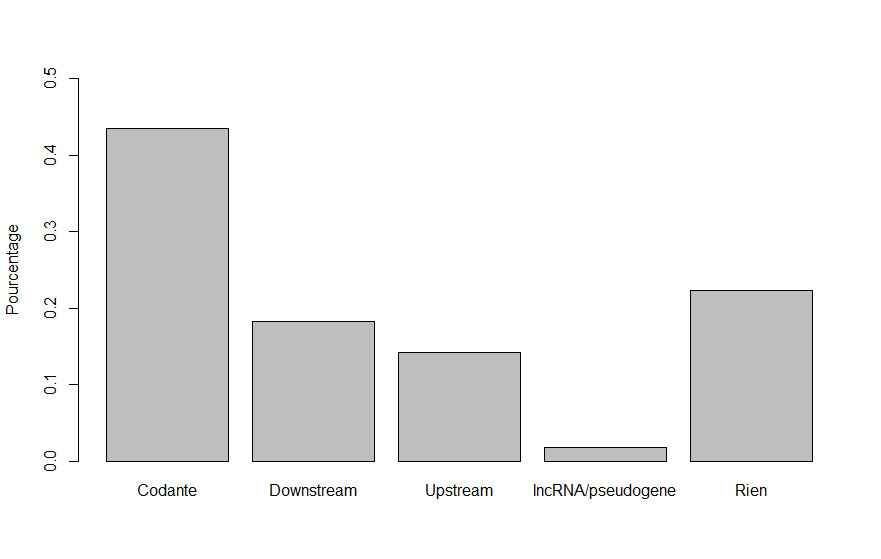

**Figure S5**

“Pourcentage” = Percentage

(A) **Molecular functions**

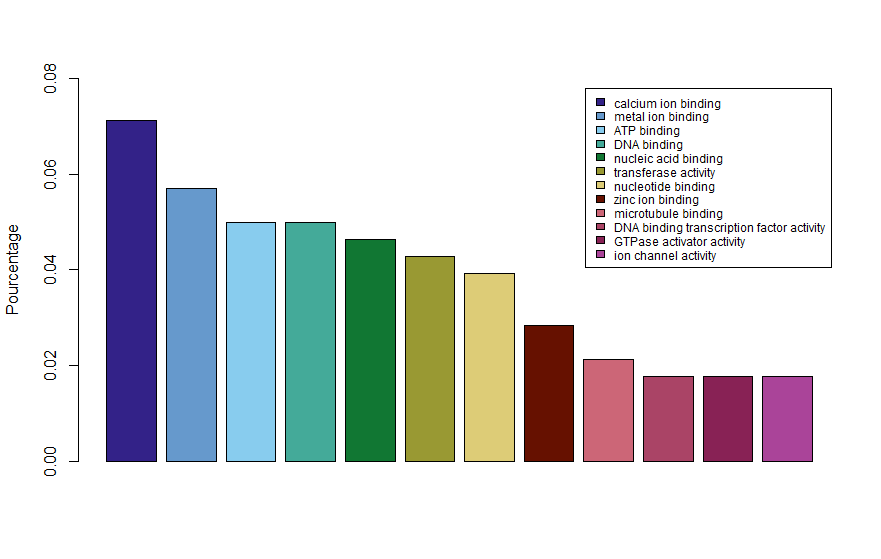

(B) **Biological processes**

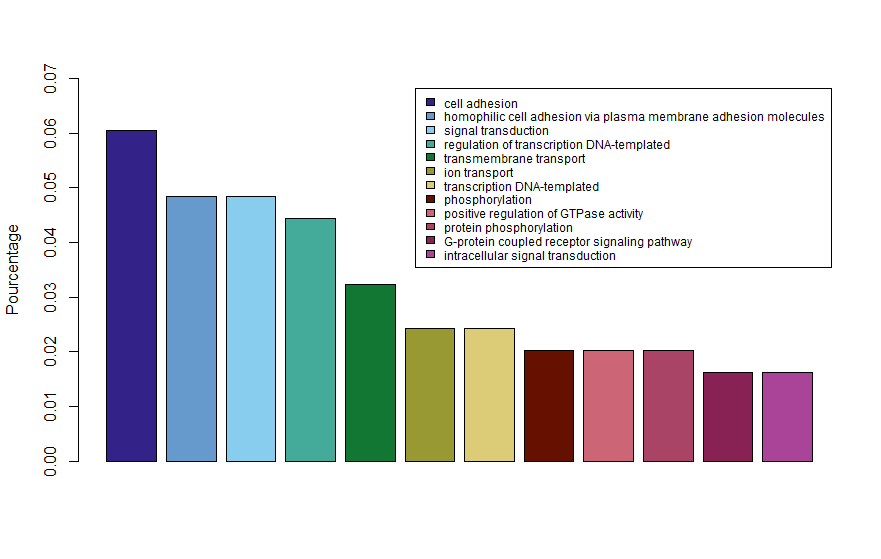
